# Genes derived from ancient polyploidy have higher genetic diversity and are associated with domestication in *Brassica rapa*

**DOI:** 10.1101/842351

**Authors:** Xinshuai Qi, Hong An, Tara E. Hall, Chenlu Di, Paul D. Blischak, Michael T. W. McKibben, Yue Hao, Gavin C. Conant, J. Chris Pires, Michael S. Barker

## Abstract

- Many crops are polyploid or have a polyploid ancestry. Recent phylogenetic analyses have found that polyploidy often preceded the domestication of crop plants. One explanation for this observation is that increased genetic diversity following polyploidy may have been important during the strong artificial selection that occurs during domestication.
- To test the connection between domestication and polyploidy, we identified and examined candidate genes associated with the domestication of the diverse crop varieties of *Brassica rapa*. Like all “diploid” flowering plants, *B. rapa* has a diploidized paleopolyploid genome and experienced many rounds of whole genome duplication (WGD). We analyzed transcriptome data of more than 100 cultivated *B. rapa* accessions.
- Using a combination of approaches, we identified more than 3,000 candidate genes associated with the domestication of four major *B. rapa* crop varieties. Consistent with our expectation, we found that the candidate genes were significantly enriched with genes derived from the Brassiceae mesohexaploidy. We also observed that paleologs were significantly more diverse than non-paleologs.
- Our analyses find evidence for that genetic diversity derived from ancient polyploidy played a key role in the domestication of *B. rapa* and provide support for its importance in the success of modern agriculture.

## Introduction

Polyploidy, or whole genome duplication (WGD), has long been associated with crop domestication and diversity (Anderson, 1969; Lewis, 1980; Heiser, 1990; Hilu, 1993; Paterson, 2005; Udall & Wendel, 2006; Meyer *et al*., 2012; Renny-Byfield & Wendel, 2014). Many desirable crop traits such as larger seed size, greater stress tolerance, and increased disease resistance are often attributed to polyploidy (Lewis, 1980; Levin, 1983). A recent phylogenetic analysis found that domesticated plants have experienced significantly more polyploidy than their wild relatives (Salman-Minkov *et al*., 2016). Polyploidy often precedes domestication and crops are nearly twice as likely to be domesticated in lineages with a relatively recent WGD compared to those without (Salman-Minkov *et al*., 2016). Among the potential explanations for the relationship between polyploidy and domestication, the expanded genetic diversity and plasticity of polyploid plants may be especially advantageous during domestication and crop improvement (Otto & Whitton, 2000; Shimizu-Inatsugi *et al*., 2017; Baduel *et al*., 2018; Paape *et al*., 2018; Monnahan *et al*., 2019). Analyses in yeast have shown that polyploid lineages not only have higher genetic diversity but also adapt to new environments faster than their lower ploidal level relatives (Selmecki *et al*., 2015). Similarly, the niches of polyploid plants evolve faster than their diploid relatives (Baniaga *et al.,* 2020). These features may collectively give polyploids unique advantages over diploids during domestication and the global spread of crops that occurred with human population expansion.

Although nearly 30% of plant species are recent polyploids, all flowering plants are paleopolyploids with varying histories of WGD (One Thousand Plant Transcriptomes Initiative, 2019). Given that the genetic consequences of polyploidy play out over time as genomes diploidize and paralogs fractionate (Otto, 2007; Arrigo & Barker, 2012; Baduel *et al*., 2018), we may expect that the effects of polyploidy extend to diploidized species. Here, we sought to test whether past polyploidy is associated with increased diversity and domestication in the crop varieties of *Brassica rapa*. Like all flowering plants, the genome of *B. rapa* has been multiplied and fractionated many times over. The most recent polyploidization in the ancestry of *B. rapa* was a mesohexaploidy that occurred approximately 9–28 MYA(Lukens *et al*., 2004; Lysak et al., 2005; Beilstein et al., 2010; Wang et al., 2011; Arias et al., 2014; Cheng *et al*., 2014; Franzke *et al*., 2016). Further, *B. rapa* has been domesticated into many different crop varieties across Europe and Asia. These include turnips, oil seeds, pak choi, Chinese cabbage, and mustard seeds. Many researchers have suggested that there is a connection between the mesohexaploidy and the diversity of *B. rapa* crop varieties (Cheng *et al*., 2014, 2016), but the relationship has never been explicitly tested. Using recently sequenced transcriptomes from a diverse array of *B. rapa* accessions (Qi *et al*., 2017), we tested if polyploid-derived regions of the genome are enriched with candidate genes associated with domestication. We also compared genetic variation in the polyploid vs non-polyploid derived regions of the *B. rapa* genome. Given the frequency of ancient polyploidy and its contribution to the evolution of plants, our analyses demonstrate the key role of polyploidy in the domestication of *B. rapa* and provide support for its importance in the success of modern agriculture.

## MATERIALS & METHODS

### Data Sources

RNA-seq data for this study were previously generated by our group (Qi *et al*., 2017). Based on previous population genomics analyses (Qi et al 2017), a total of 102 *B. rapa* accessions (Supplementary Table 2) were selected from the USDA GRIN database (http://www.ars-grin.gov/) or from the author’s collection to represent the five major *B. rapa* genetic groups. The five genetic groups include an earlier derived Europe and Central Asia group, represented by turnip (TN, *B. rapa* and *B. rapa* subsp. *rapa,* 22 accessions); four derived *B. rapa* groups, represented by Pak choi (PC, *B. rapa* subsp. *chinensis,* 25 accessions), Chinese cabbage (CC, *B. rapa* subsp. *pekinensis,* 28 accessions), Indian sarson (IS, *B. rapa* subsp. *trilocularis* and *B. rapa* subsp. *dichotoma*; 20 accessions) and toria (TO, 7 accessions). The last four genetic groups diverged from the TN group about 2400–4100 years ago after the initial *B. rapa* domestication in European-Central Asia (Qi et al., 2017). Two *B. oleracea* accessions (SRR630924 and SRR1032050) from the NCBI Sequence Read Archive (SRA) were used as outgroups for analyses.

### RNA-seq Variant Calling

Raw reads were cleaned using Trimmomatic version 0.32 (Bolger *et al*., 2014), and mapped to the reference *B. rapa* genome (version 1.5, http://brassicadb.org/brad/) with Tophat version 2.0.14 (Trapnell *et al*., 2009). SNP calling was performed using Samtools version 0.1.18 and bcftools version 0.1.17 *(Li et al., 2009; Li, 2011)*. The resulting VCF files were filtered with the vcfutils.pl script and vcffilter. Only SNPs with depth greater than 10 and variant quality (QUAL) greater than 30 were retained, which included 1.32 million SNPs. Details about sequencing and variant calling were described previously (Qi *et al*., 2017).

### Identifying Genes Derived from the Mesohexaploidy Event (Paleologs)

We used two different approaches to identify paleologs—genes derived from an ancient polyploid event—retained from the Brassiceae mesohexaploidy, the most recent polyploidy in the ancestry of *B. rapa.* Both approaches aim to distinguish genes that were born from the mesohexaploidy from genes derived from other types of duplication events, such as tandem, segmental, and other small-scale duplications. Genes that we identify as derived from the Brassiceae mesohexaploidy were labelled as paleologs, and other genes were labelled as non-paleologs. The first approach used synteny and gene age distribution data from only *B. rapa* to infer paleologs. To estimate the age distribution of gene duplications (also called a Ks plot) for *B. rapa* we obtained CDS sequences from the *Brassica* database (http://brassicadb.org/brad/). The synonymous substitution rate (Ks) of each gene duplication was calculated using DupPipe (Barker *et al*., 2008, 2010). The gene age distribution was visualized in R using histograms. The boundaries of the hexaploid peak were determined using EMMIX (McLachlan *et al*., 1999) by fitting a mixture model of normal distributions to the Ks data (Tiley *et al*., 2018). One hundred random starting points and 10 k-means starting points were used to identify the number of normal distributions (from 1 to 10). The best fit model was selected based on the Bayesian information criterion (BIC) value. Ks nodes with >50% likelihood assignment to the mesohexaploidy peak were considered as gene pairs derived from the *Brassica* hexaploidy event. We combined these age based inferences of paleologs with syntenic data for *B. rapa*. Syntenic gene sets were generated by CoGe SynMap (genomevolution.org/CoGe/SynMap.pl) with a 2: 2 quota-align ratio and default parameters (Tang *et al*., 2011) using the unmasked *B. rapa* genome (Version 1.5). These two paleolog lists were integrated with an independently estimated *B. rapa* paleolog list (Cheng *et al*., 2012). Genes present in at least two of these three lists were extracted using the Venn online tool (http://bioinformatics.psb.ugent.be/webtools/Venn/) and were considered to be high confidence paleologs. Genes only present in one of these three lists were considered low confidence paleologs and were not used in our study. Genes not present in any of the above three paleologs lists were considered non-paleologs.

We also developed a second set of paleologs and non-palelogs from a recent multi-genome comparative analysis with POInT, the Polyploidy Orthology Inference Tool (Hao *et al*., 2020). The POInT analysis modeled the history of genome evolution following the Brassiceae mesohexaploidy by comparing the genomes of *Arabidopsis thaliana*, *Brassica oleracea*, *Brassica rapa*, *Crambe hispanica*, and *Sinapis alba (REF)*. Briefly, the POInT analysis compared each of the Brassiceae species genomes to *A. thaliana* to identify blocks of triple conserved synteny across each of the four genomes. Blocks from each genome were then merged into “pillars” that contain one to three surviving paralogs in each species from the mesohexaploidy. POInT uses a probabilistic approach to assess if a gene belongs to a paleolog pillar. We filtered for genes in the *B. rapa* genome that had a >95% probability of assignment to one of these mesohexaploid pillars from the Hao et al. (2020) analysis. These data were used to develop a list of *B. rapa* paleologs that were retained in single, double, or triple copy from the ancient hexaploidy. All other genes in *B. rapa* were then classified as non-paleologs.

### Selection analyses

We used two different methods to identify genes with signals of positive selection in the four derived *B. rapa* genetic groups. Genomic regions with evidence consistent with selective sweeps were detected using SweeD 3.0 *(Pavlidis et al., 2013)* based on the composite likelihood ratio (CLR) test of SNP site frequency spectrum (SFS) patterns. A total of 1.32 million SNPs were used in the SweeD tests for the TN group and each of the four derived *B. rapa* genetic groups. These analyses were performed with the default settings except each chromosome was divided to 60,000 grids. The CLR was calculated for each equally generated relative position on each chromosome. Only the top 1% significant outlier regions were considered. Genes within the outlier regions were then annotated based on *B. rapa* genome v1.5.

We also used the McDonald-Kreitman test (M-K test) (McDonald & Kreitman, 1991) to identify genes experiencing positive selection. This method is based on the proportion of synonymous and nonsynonymous substitutions within and between species that are due to directional selection. For the M-K test, we compared SNPs specific to each of the four *B. rapa* crop groups with the two *B. oleracea* outgroup accessions. For each separate SNP dataset, we annotated synonymous and nonsynonymous SNPs using SnpEff v4.2 (Cingolani *et al*., 2012). The reference genome database was created using *B. rapa* genome v1.5. The significance of the M-K test was evaluated using Fisher’s exact test.

### Differential gene expression analysis

Differential gene expression analysis was performed following the Tuxedo protocol (Trapnell *et al*., 2012). Briefly, cleaned reads were first mapped to the *B. rapa* genome with TopHat (Trapnell *et al*., 2009), then mapped BAM files were assembled in Cufflinks (Trapnell *et al*., 2010), merged with Cuffmerge, quantified in Cuffdiff (Trapnell *et al*., 2013) (Cufflinks version2.2.1) with multiple-testing corrected q-values as 0.05, and finally indexed and visualized in CummeRbund version 2.14 (Goff *et al*., 2012). Genes significantly differentially expressed between the TN group and the four derived *B. rapa* groups were identified using the getSig function in CummeRbund (Goff *et al.,* 2012).

### Candidate genes from the literature

To confirm our findings, we compared our observations of paleolog enrichment with a list of candidate genes from published studies of *B. rapa*. We surveyed the literature for fine mapping and bulk segregant analyses of *B. rapa* that mapped traits to one or a few candidate genes. These studies and the candidate genes are listed in Supplemental Table 7. Overall, our survey identified 40 candidate genes in *B. rapa* that have been mapped to the location of classical loci associated with crop traits.

### Statistical analyses

To test for evidence of paleolog enrichment in our candidate gene lists, we used 2X2 chi-square tests. We compared the numbers of paleologs and non-paleologs that were not candidate genes to the numbers of paleolog and non-paleolog candidate genes. For each candidate gene list, we aggregated the candidate genes of the four derived *B. rapa* genetic groups. The ratio of paleologs and non-paleologs among non-candidate genes in the *B. rapa* genome was used as the expected value for the chi-square test to determine if the candidate gene lists contained more paleologs than expected.

To demonstrate the logical relations among these lists, Venn diagrams were generated using the online Venn diagram tool (http://bioinformatics.psb.ugent.be/webtools/Venn/). The density distribution of the identified paleologs, non-paleologs, SweeD outlier gene, M-K test outlier gene and differentially expressed gene were visualized in Circos with a bin size of 100 kbp. We used the dhyper function in R to test for significant overlap among the different candidate gene lists.

### Nucleotide diversity of *B. rapa* genes

The nucleotide diversity (*π*) of each paleolog and non-paleolog gene was estimated using a custom script from the output of VCFtools (Danecek *et al*., 2011). The results were summarized and visualized in R v3.6.1 (R Core Team, 2019).

As a secondary verification of our calculations for nucleotide diversity in paleologs versus non-paleologs, we remapped the filtered RNAseq reads to the *B. rapa* reference genome using STAR v2.7.3a (Dobin *et al*., 2013). We then used the resulting BAM files to calculate nucleotide diversity using genotype likelihoods with ANGSD v0.921 (Korneliussen *et al*., 2014). This procedure was done twice, once filtering for uniquely mapping reads and once without, to see if the results differed based on potential read mismapping caused by gene duplicates. For these calculations, we first inferred the folded site frequency spectrum (-doSaf 1, -fold 1) and used this as a point estimate for estimating theta (-doThetas 1) at each site separately for all paleologs and nonpaleologs (Nielsen *et al*., 2012; Korneliussen *et al*., 2013). We then used the thetaStat program within ANGSD to calculate nucleotide diversity for each gene. We also compared the distributions of nucleotide diversity between paleologs and nonpaleologs using both t-tests and non-parametric Mann-Whitney U-tests within R v3.6.1 (R Core Team, 2019).

## Results

### Partitioning the *B. rapa* genome into paleologs vs non-paleologs

To test the contribution of paleopolyploidy to the domestication of *Brassica rapa,* we used two different approaches to classify genes as paleologs—genes derived from the *Brassica* mesohexaploidy—or non-paleologs. In the first approach, we used gene age distributions and synteny in *B. rapa*. An initial list of putative paleologs was generated from pairs of genes with synonymous divergence (Ks) values that correspond to the peak of duplication associated with the Brassiceae mesohexaploidy. This approach identified 21,280 genes that are likely derived from the mesohexaploidy (Fig. 1a). This result is consistent with previous *Ks* estimates (Barker *et al*., 2009; Cheng *et al*., 2013). Ancient WGDs and their associated paralogs may also be classified by identifying syntenic blocks of duplication. Syntenic analyses with CoGe recovered 19,810 syntenic gene sets that contained 31,796 paleologs. Finally, we compared these two lists of putative paleologs with a previously curated list of 23,716 paleologs reported by an independent research group (Cheng *et al*., 2012). Genes that appeared at least twice in these three lists were classified as paleologs and were used in further analyses (Fig. 1b). The final paleolog list included 27,919 genes (Supplementary Table 1), which represents 68.06% of the 41,020 annotated genes in the *B. rapa* reference genome (Version 1.5). We also considered the 5,424 genes that were not present in any of the above three paleolog lists as non-paleologs (Fig. 1b), which represents 13.22% of the total *B. rapa* genes. Paleologs and non-paleologs are distributed throughout the *B. rapa* genome (Fig. 2).

**Fig. 1:**
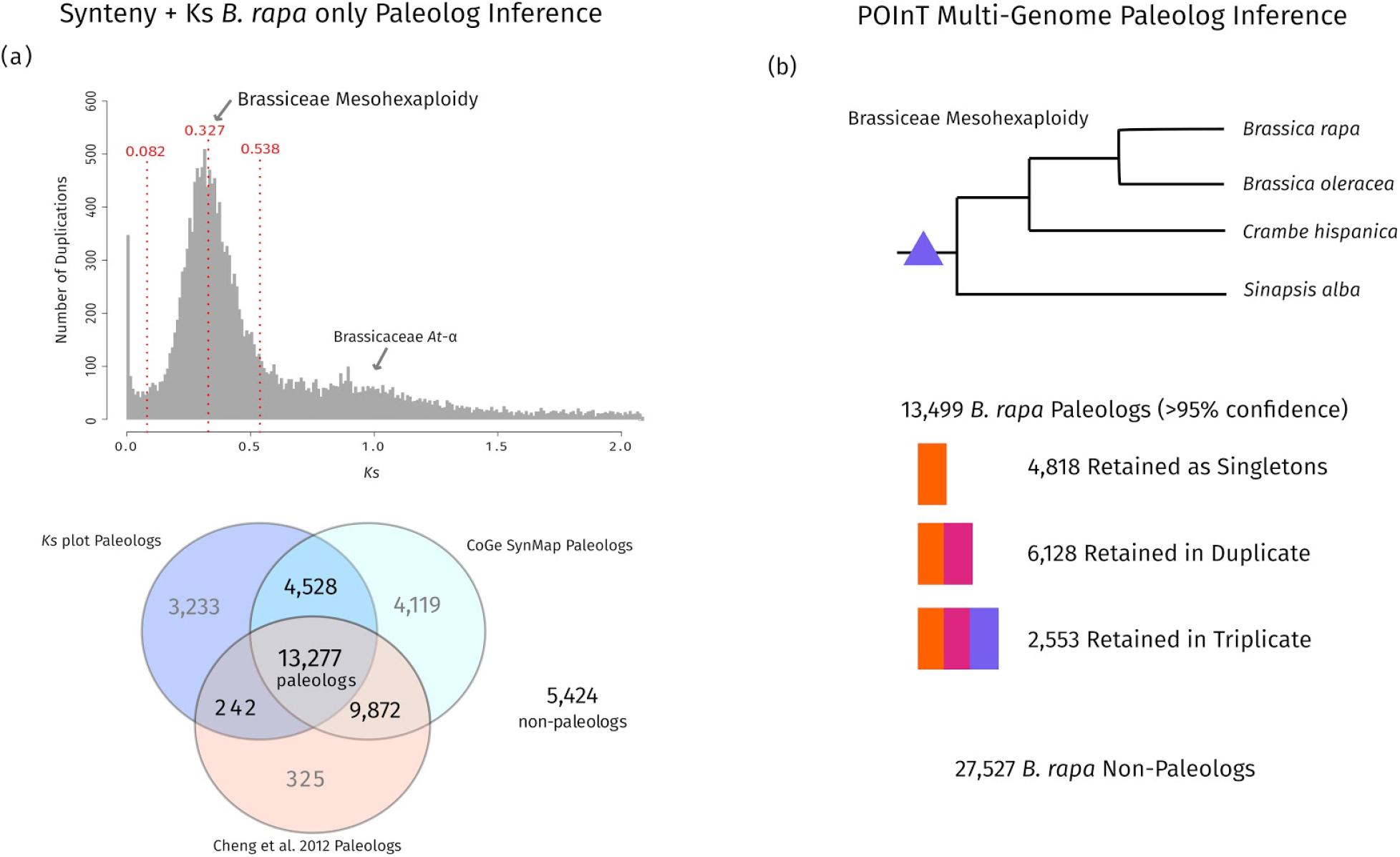
Identifying paleologs derived from the *Brassica* hexaploidy event. **(a)** The age distribution of gene duplications from 41,020 CDSs in the *B. rapa* genome (version 1.5). The x-axis represents the synonymous divergence of duplication events (Ks value), whereas the y-axis represents the number of duplications. Gene pairs with Ks divergence 0.082–0.538 were identified as putative paleologs. Venn diagram showing the overlap among our two *B. rapa* paleolog lists with a previously reported *B. rapa* paleolog list. Genes that appeared at least twice in these three lists were considered high confidence paleologs and were used in further analyses. Genes that were not present in any of the three paleologs lists were considered to be non-paleologs. **(b)** Phylogeny of species’s genomes used for the POInT multi-genome analysis to identify paleologs in *B. rapa* from Hao et al. (2020). A total of 13,499 paleologs and 27,527 non-paleologs were identified in *B. rapa*. Colored blocks indicate the retention category of paleologs and the number of genes in each category.

**Fig. 2:**
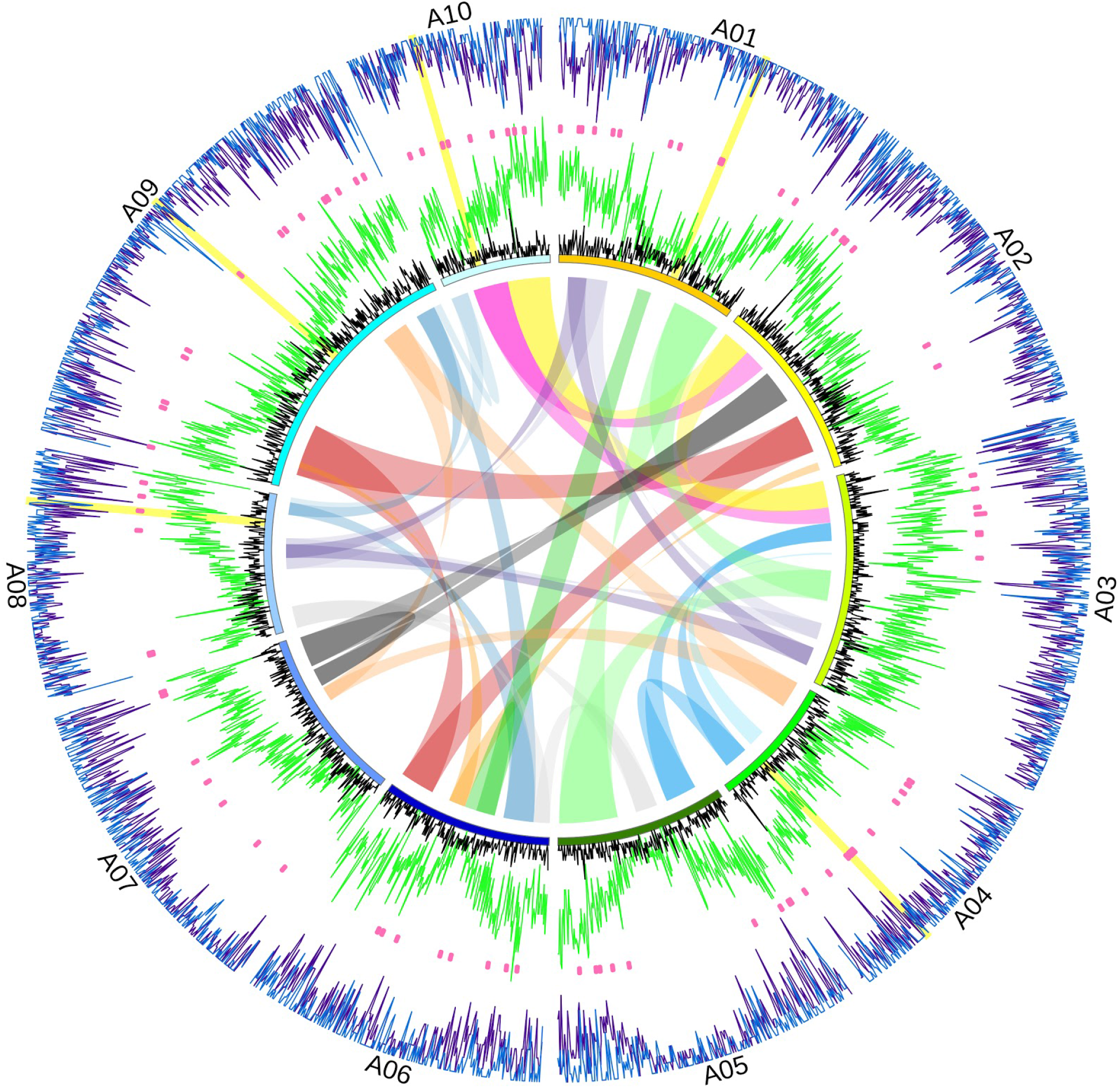
Circos plot of the distribution of surveyed genes in this study. Highlighted on the outside of the plot are the *Brassica* hexaploidy paleologs identified by the *B. rapa* only synteny + Ks analysis (light green lines), non-paleologs (black lines), and candidate genes identified by SweeD (blue lines), the McDonald-Kreitman test (pink dots), and the significantly differentially expressed genes between the EU-CA group and the four derived *B. rapa* groups (purple lines). Heights of the lines indicate the relative density of genes in each category in the sliding window at each location. A01-A10 represent the 10 chromosomes of *B. rapa*. The rainbow ribbons in the center represent the syntenic regions among chromosomes. The yellow bars represent the location of the five *B. rapa* domestication candidate genes listed in Table S5.

We also used results from a recent multi-genome comparative analysis *(Hao et al., 2020)* to infer paleologs in *B. rapa*. A recent POInT analysis of the genomes for four Brassiceae species *(Brassica oleracea*, *Brassica rapa, Crambe hispanica*, and *Sinapis alba)* uncovered evidence for 14,050 regions (“pillars” in POInT terminology) of triple conserved synteny from the Brassiceae mesohexaploidy across the four genomes. Genes in these regions of conserved synteny are present in one to three copies in each genome. After parsing these output for *B. rapa* genes that were assigned to these regions with >95% probability by POInT, we identified 4,818 paleologs retained as single copy genes, 6,128 paleologs retained in duplicate, and 2,553 retained in triplicate (FIG 1b; Supplementary Table 1). The remaining 27,527 genes in the *B. rapa* genome were considered non-paleologs. Using this approach, only 32.9% of the *B. rapa* genome were identified as paleologs, a much lower fraction than the above synteny + Ks *B. rapa* only method. Despite the difference in the number of inferred paleologs, there was a high overlap in the lists with 89.2% of the paleologs inferred by POInT also found by the *B. rapa* only approach. Most of the difference in the overall number is likely due to the POInT approach’s initial filtering for regions of the genome that have been retained in triplicate and requiring that each of these regions is found across all species, greatly reducing the number of paleologs identified. In contrast, the B. rapa only approach recovers more paleologs because it only requires regions to be retained in duplicate in the genome of only one species (*B. rapa).* Notably, a less stringent version of the POInT results (not presented) that included all possible identified paleologs regardless of assignment probability identified 22,774 paleologs with 89.5% overlap with the *B. rapa* only list. Thus, these methods appear to be largely identifying the same genes as paleologs in the *B. rapa* genome. These methods provide two different assessments of paleolog identity that we used to further explore the relationship between polyploidy and domestication of *B. rapa*.

### Analyses of selection identify candidate genes associated with domestication of *B. rapa* crop varieties

We used a diverse collection of sequenced transcriptome data from across five groups to identify candidate genes associated with the domestication and improvement of *B. rapa* crop varieties. In total, 1.32 million SNPs from transcriptome data with 25X coverage or greater from 102 *B. rapa* accessions were analyzed (Supplementary Table 2). Based on our previous population genomic analyses of these data (Qi *et al*., 2017), the accessions are comprised of a European-Central Asian *B. rapa* population represented by turnip (TN, *B. rapa* subsp. *rapa)* and four derived groups represented by pak choi (PC, *B. rapa* subsp. *chinensis),* Chinese cabbage (CC, *B. rapa* subsp. *pekinensis),* Indian sarson (IS, *B. rapa* subsp. *trilocularis)* and toria (TO, *B. rapa* subsp. *dichotoma).* We analyzed these data with an ensemble of molecular evolution and population genomic approaches to identify candidate genes associated with each group.

During domestication and crop improvement, we expect genetic variation important for agricultural traits to be selected in crop populations. To identify genes or genomic regions that have experienced recent positive selection, we used two different approaches. First, we used a selective sweep test, SweeD (Pavlidis *et al*., 2013), to identify regions associated with significantly reduced variation consistent with a recent selective sweep in each of the crop varieties. We identified 3,387 unique genes within the identified selective sweep regions (Fig. 3a, Supplementary Tables 3 and 4, and Supplementary Fig. 1 and 2). This included 1,072 genes in *chinensis,* 687 genes in *pekinensis,* 1,048 genes in toria, 1,228 genes in the sarsons, and 1,713 genes in European-Central Asian group. On average 70% of these genes were found in only one crop variety (Fig. 3a), indicating that many of these genes may have swept during the putative independent domestication of each crop variety. We also used the McDonald-Kreitman test (M-K test) (McDonald & Kreitman, 1991) to identify coding regions with a significant excess of fixed amino acid substitutions, a different signature of positive selection. We used *B. oleracea* as an outgroup. Analyzing each crop variety with the M-K test, we found 92 genes in total with a molecular evolutionary signature of positive selection (Fig. 3b and Supplementary Tables 3 and 4). All but one of these genes was uniquely identified in a different crop variety, similar to the selective sweep analysis and consistent with independent domestication and differentiation of these *B. rapa* crops. Overall, we identified 3,479 candidate genes that may be associated with the domestication and improvement of the five *B. rapa* crop groups.

**Fig. 3:**
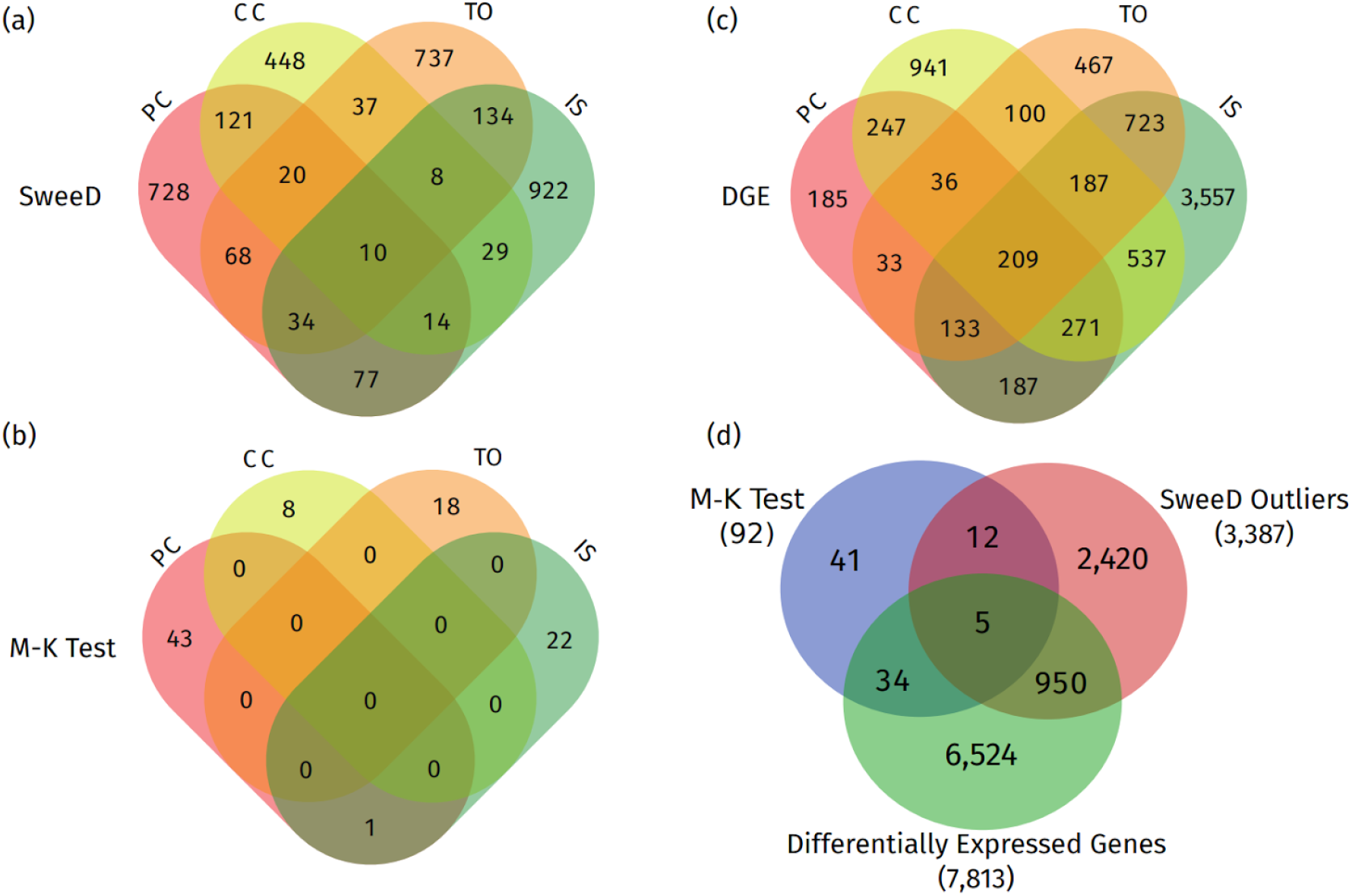
Overlap among domestication candidate genes. PC = pak choi, CC = Chinese cabbage, TO = toria, and IS = Indian sarson. Number and overlap of candidate genes for each crop variety inferred by **(a)** SweeD, **(b)** the McDonald-Kreitman test, and **(c)** differential gene expression analyses. **(d)** Total number and overlap of candidate genes inferred by different methods across all crop varieties.

We may also expect significant changes in gene expression to occur during domestication and crop improvement that may not be apparent in analyses of positive selection. To identify candidate genes with changes in gene expression during domestication, we performed differential gene expression (DGE) analyses with the four derived *B. rapa* genetic groups. RNA-seq data for each individual was collected 20 days after germination and the expression levels of genes in each group were compared to gene expression levels in the TN group. We identified a total of 7,813 genes with significant differential expression when comparing each group with the TN population (Fig. 3c, Supplementary Tables 3 and 4, and Supplementary Fig. 2 and 3). Unlike the analyses of selection above, the number of unique differentially expressed genes varied widely among the different crop varieties. The sarsons (IS) were the most differentiated from TN and the other crops with over 61% of their differentially expressed genes unique. The sarsons also had the largest number of differentially expressed genes at 5,804. In contrast, pak choi (PC) had the smallest number of differentially expressed genes, 1,301, and the lowest percentage that were uniquely different at 14.2%. Other crop varieties fell between these values. The genes identified by all three methods were largely different among all the crops (Fig. 3d) with only a small number of genes identified by one or more of these approaches (i.e., sweep test, M-K test, or DGE analyses). However, the number of overlapping genes found by each of these approaches was significant in all cases based on the expected numbers from a hypergeometric distribution (MK:SweeD overlap *p* = 0.0013; MK:DGE overlap *p* < 0.0001; SweeD:DGE overlap *p* < 0.0001). This suggests that we are detecting shared signatures of domestication across these different candidate gene approaches. Similarly, most of the genes found to be associated with recent positive selection or differential gene expression were unique to a particular crop lineage as expected with independent domestication and differentiation of these distinct crop varieties.

Across all of the analyses, only five genes were repeatedly present in tests among one or more of the crop varieties (Fig. 3d and Supplementary Table 5). These genes—*Bra010933, Bra023190*, *Bra031452, Bra003055,* and *Bra035693—were* identified in some combination of the three inference methods with evidence for both signatures of positive selection and changes in gene expression. Considering that we recovered these five genes using different approaches, they may play an important role during the differentiation and improvement of *B. rapa* crop varieties. Searches of these genes and their *Arabidopsis* homologs in the STRING (Szklarczyk *et al*., 2019) and UniProt (UniProt Consortium, 2018) databases recovered a diverse range of potential functions. *Bra023190* i s uncharacterized in *B. rapa,* but is homologous with *SGR2* in *Arabidopsis thaliana,* a gene associated with negative gravitropism and leaf movement in darkness (Kato *et al*., 2002; Mano *et al*., 2006). Notably, this gene was identified as a candidate gene in both of the cabbage crops analyzed here, *B. rapa chinensis* and *pekinensis*. Other genes are homologs with *A. thaliana* genes implicated in responses to oxidative stress from photooxidation *(Bra010933),* salt stress *(Bra031452*; (Tuteja *et al*., 2012)), and a vacuolar V-type proton ATPase *(Bra035693).* Finally, *Bra003055,* identified by all three candidate gene approaches in the sarsons, was previously found to be over-expressed in *B. rapa* in soils that are deficient in iron and with an excess of zinc (Li *et al*., 2014).

### Paleologs from the *Brassica* mesohexaploidy and domestication

To test if the candidate genes associated with domestication of the *B. rapa* crop varieties are enriched with paleologs, we compared the number of paleologs in the candidate gene lists to the expected number from our two genome-wide surveys (Fig. 1). We found that the candidate genes were significantly enriched with paleologs from the Brassiceae mesohexaploidy using either paleolog inference method (Fig. 4). Using paleologs inferred with the *B. rapa* only synteny + Ks approach (Fig. 1a), genes from all three candidate gene approaches were enriched across all crops with paleologs comprising 78.54% of SweeD (*χ*^2^ = 160.06, *p* < 0.001), 83.7% of McDonald Kreitman (*χ*^2^ = 10.33, *p* = 0.0013), and 78.6% of differentially expressed candidate genes (*χ*^2^ = 345.40, *p* < 0.001). Similarly, we found that candidate genes were significantly enriched with paleologs inferred with the POInT multi-genome approach (Fig. 1b). Using the POInT defined paleologs, we found that 42.6% of SweeD (*χ*^2^ = 157.84, *p* < 0.001), 53.2% of McDonald Kreitman (*χ*^2^ = 16.40, *p* < 0.001), and 40.3% of the differentially expressed candidate genes (*χ*^2^ = 239.04, *p* < 0.001) were paleologs. The POInT method allowed us to identify genes retained in single, double, and triple copy from the Brassiceae mesohexaploidy and we found that these varied in their enrichment among the candidate gene lists (Fig. 4; Supplementary Table 6). Paleologs that have returned to single copy in *B. rapa* were significantly enriched in all of the candidate gene lists, whereas paleologs that are retained in duplicate were enriched in the SweeD and differentially expressed gene lists. In contrast, paleologs that have retained all of their copies since the mesohexaploidy—those that are retained in triplicate—were only found to be significantly enriched in the differentially expressed gene candidate gene list, but not in either candidate gene list where there is evidence of significant protein coding changes among the genes. Overall, these results suggest that genes derived from the Brassiceae mesohexaploidy were preferentially selected during the domestication of the *B. rapa* crops.

**Fig. 4:**
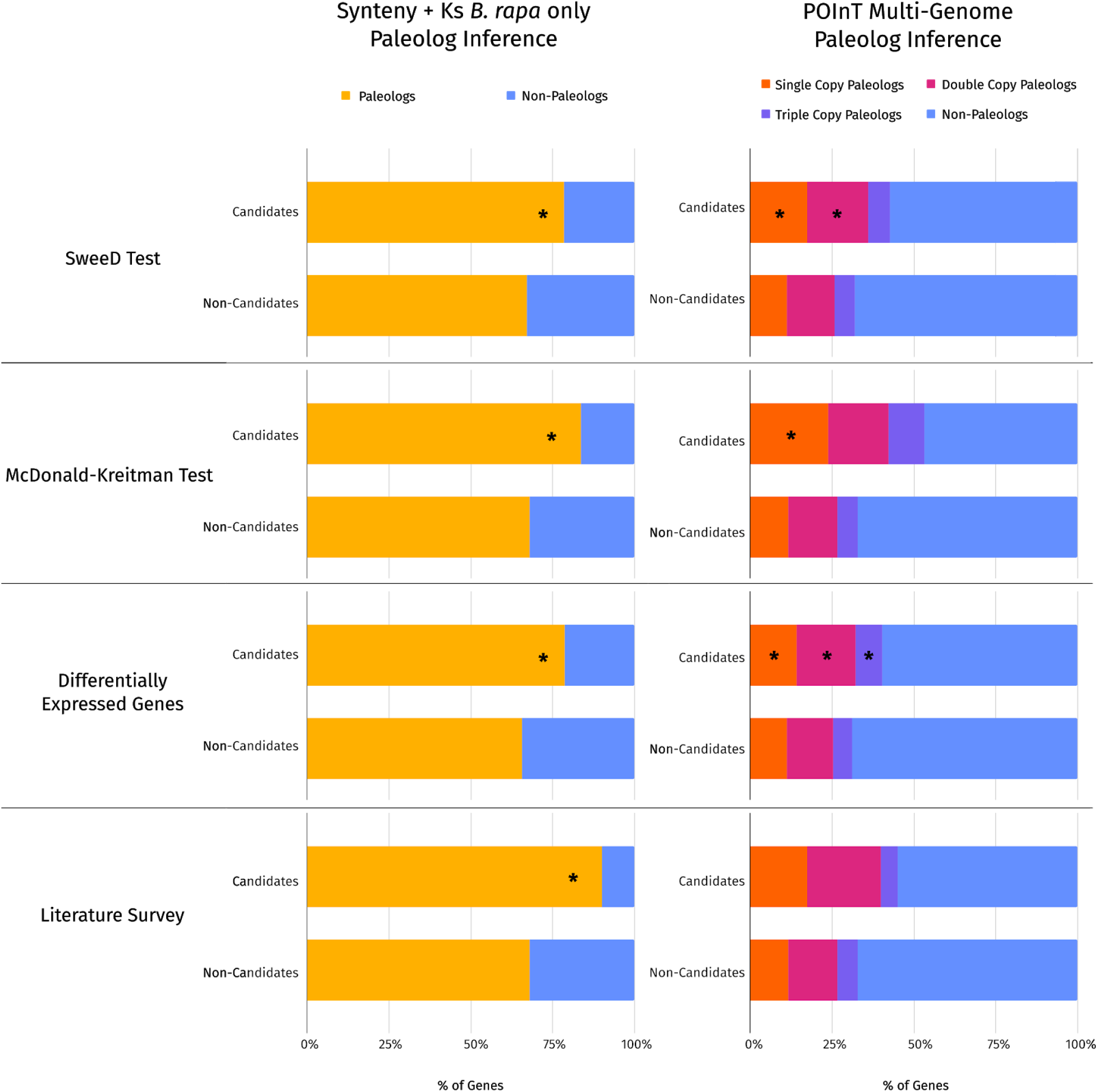
Comparison of paleolog percentages across candidate gene lists. Percentage of paleologs in each candidate gene list compared to the rest of the *B. rapa* genome. Bar charts on the left are the paleologs (yellow) and non-paleologs (blue) identified by the *B. rapa* only synteny + Ks paleolog inference approach. Bar charts on the right are paleologs identified by the POInT multi-genome approach and include paleologs retained in single (orange), double (magenta), and triple (purple) copy as well as the percentage of the *B. rapa* genome that are non-paleologs (blue). The asterisks indicate significant deviation from the null based on chi-square tests *(p* < 0.05).

To further test if domestication genes in *B. rapa* are significantly enriched with paleologs, we also developed a list of candidate genes from the literature. We focused on studies published over the last 10 years that identified genes through other approaches, such as fine mapping or bulk segregant analysis, to better establish a causal relationship between loci and crop traits. In total, we identified 40 candidate genes that fit these criteria from the literature (Supplementary Table 5). Many of these genes are associated with leaf and seed color variation, clubroot resistance, and cuticular wax biosynthesis. Notably, 15 of these genes were recovered in our candidate gene scans (Supplementary Table 7). Four of these genes were identified in our selective sweep and differential gene expression analyses. Mapping studies previously identified these genes as being associated with leaf color variation (Bra006208)*(Fu et al., 2019)*, cuticular wax biosynthesis (Bra011470 and Bra032670; (Wang *et al*., 2017, 2019), and clubroot resistance (Bra019410; (Yu *et al*., 2016). For other loci, mapping studies have identified a small collection of candidate genes in target regions. For example, *Rcr5* is a gene of major effect for clubroot resistance in *Brassica rapa*. A recent bulk segregant analysis and fine mapping study found eight genes in the *Rcr5* target region of *B. rapa (Huang et al., 2019*). In our analyses, three genes in the center of their region were found to be significantly differentially expressed. These results suggest that future mapping studies in *B. rapa* may be able to leverage our candidate gene lists to improve gene identification.

How many of the candidate genes from our literature survey were paleologs? Of the 40 genes identified in the literature, 36 were paleologs with the *B. rapa* synteny + Ks method and 18 were paleologs with the POInT approach (Fig. 4). The literature candidate genes were significantly enriched with paleologs defined by the *B. rapa* only approach (*χ*^2^ = 8.85, *p* = 0.0029) similar to our three different candidate gene lists. Although the percentage of POInT paleologs was higher in the literature candidate gene list than the rest of the genome (Fig. 4), it was not a significant difference (*χ*^2^ = 2.13, *p* = 0.1441). Given these literature-based candidate genes were largely identified with mapping based approaches rather than genomic scans, these results suggest that the paleolog enrichment observed in our analyses is not likely an artifact of the inference methods. Overall, our results indicate that paleologs were an important source of variation for domestication in *B. rapa*.

Why did artificial selection preferentially target paleologs? One possible explanation is that these genes may harbor more genetic diversity due to their paralogous history over the past 20 million years. To test this hypothesis, we used three different approaches to examine the nucleotide diversity per gene across the *B. rapa* genome. First, we estimated nucleotide diversity in VCFtools using only uniquely mapped reads from Tophat to minimize error from incorrectly mapped reads. We found that the genes derived from the mesohexaploidy were significantly more diverse than the other genes in the genome regardless of how we identified paleologs (Fig. 5). Paleologs identified by the *B. rapa* only approach were significantly more diverse than non-paleologs (t = 43.721, df = 6535, *p* < 2.2 x 10^-16^ for t-test; W = 70,733,526 and *p* < 2.2 x 10^-16^ for U-test), and had four times the mean and an order of magnitude more median nucleotide diversity (mean *π* = 0.408 × 10^-3^, median *π* = 0.170 × 10^-3^) than the non-paleolog fraction of the genome (mean *π* = 0.111 × 10^-3^, *π* = 0.124 × 10^-4^).

**Fig. 5:**
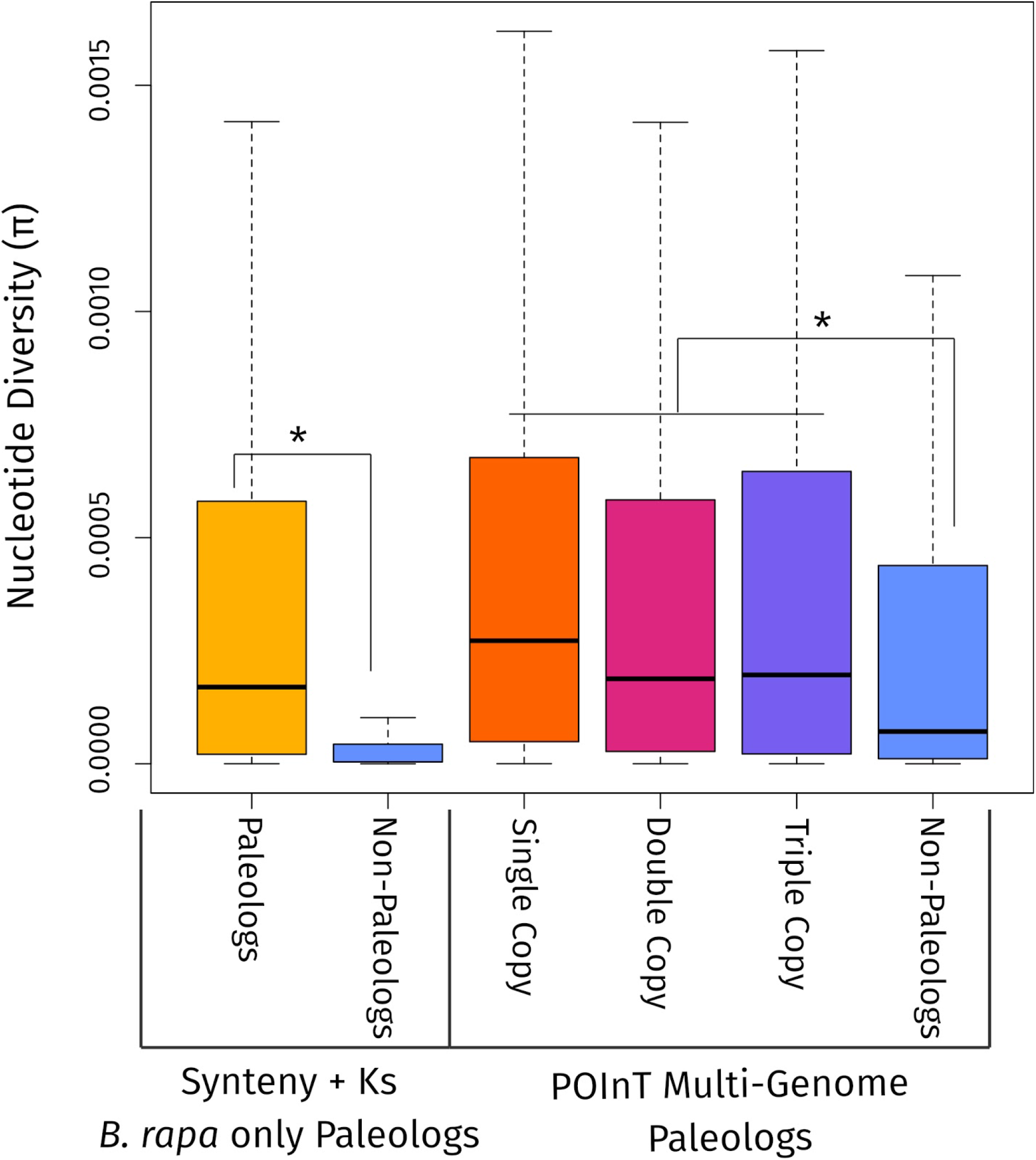
Comparison of nucleotide diversity (π) across *Brassica rapa* paleologs and non-paleologs. The bottom and top of each box represents the first and third quartiles, and the band inside each box is the median. Asterisks indicate significant differences between categories with a Mann-Whitney U test (*p* < 0.05).

This was consistent when considering only synonymous or non-synonymous substitutions (Fig. S4). POInT paleologs were also significantly more diverse than non-paleologs (t = 14.828, df = 25101, *p* < 2.2 x 10^-16^ for t-test; W = 128,146,882 and *p* < 2.2 x 10^-16^ for U-test), with a more modest difference in mean paleolog nucleotide diversity (mean *π* = 0.433 × 10^-3^, median *π* = 0.217 × 10^-3^) compared to non-paleologs (mean *π* = 0.335 × 10^-3^, median *π* = 0.713 × 10^-4^). However, the median nucleotide diversity of POInT paleologs was still an order of magnitude higher than non-paleologs. All categories of retained POInT paleologs—single, double, and triple retained copies—were significantly more diverse than the non-paleologs (Supplemental Table 6).

To further validate these results, we re-mapped reads with STAR *(Dobin et al.,* 2013) and calculated nucleotide diversity with ANGSD (Korneliussen *et al*., 2014). Although the empirical estimates of *π* were different because of differences in how *π* is calculated in these approaches, the ANGSD results recapitulated those from Tophat and VCFtools (Fig. S5). Focussing on the paleologs inferred with the *B. rapa* synteny + Ks approach, paleologs were again found to have significantly higher levels of diversity than non-paleologs (Fig. S4; t = 48.997, df = 25842, p < 2.2 x 10^-16^ for t-test; W = 238,210,946 and p < 2.2 x 10^-16^ for U-test). Importantly, these results do not differ when filtering for only uniquely mapping reads (t = 44.11, df = 26044, and p < 2.2 x 10^-16^ for t-test; W = 232,582,598 and p < 2.2 x 10^-16^ for U-test), demonstrating that our results are unlikely to be affected by read mismapping. The higher genetic diversity of paleologs in *B. rapa* may have been important for the rapid response of these plants to artificial selection during domestication.

## Discussion

Our results establish a connection between ancient polyploidy and contemporary diversity and domestication in crop varieties of *Brassica rapa*. Although polyploidy has long been hypothesized to be important for crop domestication, phylogenetic analyses have only recently confirmed that it is a key factor (Salman-Minkov *et al*., 2016). However, the mechanisms linking polyploidy and domestication itself have remained unresolved. Our results provide support for at least one genetic mechanism linking polyploidy and domestication. Paleologs, genes retained from an ancient polyploidy, were enriched in our candidate gene lists for crops of *B. rapa*. They were enriched in all three of our lists from genome scans using different statistical approaches, as well as a list developed from fine mapping and other genetic studies in the literature. In the case of *B. rapa,* the polyploid event occurred 9–28 MYA (Lukens *et al*., 2004; Lysak *et al*., 2005; Beilstein *et al*., 2010; Wang *et al*., 2011; Arias *et al*., 2014; Cheng *et al*., 2014; Franzke *et al*., 2016), long before the crop varieties were domesticated by humans. Many of these crop varieties were only domesticated in the last few thousand years (Qi *et al*., 2017), but we still find that paleologs are over-represented in a diverse range of candidate gene lists. This suggests that even ancient polyploidy may contribute to domestication and potentially adaptation long after genomes have diploidized.

We also found that paleolog retention patterns were important during domestication. Paleologs that were retained as singletons or in duplicate were enriched in candidate gene lists that involved significant changes in protein evolution and differential gene expression. In contrast, paleologs retained in triplicate from the Brassiceae mesohexaploidy were only enriched in candidate gene lists with significant differential expression changes, but not in those with significant protein changes during domestication. These results are consistent with the dosage balance hypothesis. Genes that retain all copies to maintain dosage balance are expected to be more constrained to continue making the same protein, whereas genes that are not retained to maintain dosage-balance are expected to have less constraint on their protein coding sequences (Freeling, 2009; Jiang *et al*., 2013; Conant *et al*., 2014; Blischak *et al*., 2016; Defoort *et al*., 2019). Although analyses find that some genes returning to single-copy following WGD are also highly constrained *(De Smet et al.,* 2013; Li *et al.*, 2016), our candidate gene lists were enriched with single copy paleologs across all of our analyses. This is likely because our single-copy list includes this set of “core” genes under strong constraints that returned to single-copy rapidly following WGD as well as those that returned to single-copy more gradually over millions of years. Indeed, our single-copy paleologs are enriched with the same types of GO terms as previously identified “core” single-copy genes (Hao *et al*., 2020), so our list does include the types of genes we expect to quickly return to single-copy (De Smet *et al*., 2013; Li *et al*., 2016). The gradient of paleolog retention categories following the Brassiceae mesohexaploidy—single, double, or triple retained genes—may produce more variation in the resolution of genes that would only be able to return to single copy following a WGD. The paleologs retained in triplicate, in particular, provide a strong case to test the biology of genes under apparently strong dosage balance. Overall, the patterns of paleolog retention and their roles in the domestication of *B. rapa* appear to follow the rules of the dosage-balance hypothesis.

Why are paleologs over-represented among the candidate genes for *B. rapa* domestication and crop improvement? Our results suggest that it may be because this class of genes harbors more genetic diversity than other genes in the genome. Across all of our analyses, paleologs consistently had significantly more nucleotide diversity than non-paleologs in *B. rapa*. Considering that many paleologs have been maintained in duplicate or triplicate for many generations since the mesohexaploidy that occurred 9–28 MYA, relaxed selection may yield elevated diversity at these genes as observed in other studies of paralog evolution (Lynch & Conery, 2000; Kondrashov *et al*., 2002; Conant & Wagner, 2003; Brunet *et al*., 2006; Aagaard *et al*., 2006; Scannell & Wolfe, 2008; Shan *et al*., 2009; Viaene *et al*., 2009; Innan & Kondrashov, 2010; Lee & Irish, 2011; Ascencio *et al*., 2017). Consistent with relaxed selection, we found that nucleotide diversity was elevated for both synonymous and nonsynonymous substitutions in *B. rapa* paleologs compared to non-paleologs. Notably, nucleotide diversity was higher across all categories of paleologs in our analyses, including paleologs that are now retained in single copy. This suggests that the higher diversity could be a legacy of past duplication rather than a product of ongoing masking at some paleolog loci. It may also be that non-paleologs are under more constraint than paleologs. However, we still observed a significant difference in diversity even in our POInT analysis where nearly ⅔ of the genome was classified as non-paleologs, suggesting that the difference is probably not entirely due to extraordinary constraint and purifying selection on non-paleologs. Further research is needed to better understand why the paleologs in *B. rapa* have elevated nucleotide diversity compared to the non-paleologs. The elevated diversity in the paleologs does not appear to be an artifact of read mapping error given that we recovered the same pattern with two different read mapping approaches and observed no difference when using only uniquely mapped reads. Relatively long Illumina read lengths (150 bp) and the nearly 30% synonymous divergence of paleologs from the Brassiceae mesohexaploidy may have limited read mapping errors caused by paralogy. Notably, the most recent paralogs were classified as “non-paleologs” together with other genes derived from duplication events other than polyploidy. Read mapping error rates would be expected to be the highest among these recent paralogs, but we consistently observed across our analyses that estimates of nucleotide diversity were significantly lower among non-paleologs than the paleologs. Regardless of the cause of the increased variation, it does not appear to be an artifact of read mapping and may explain why these genes are over-represented in the candidate gene lists. Increased genetic diversity is expected to be associated with greater phenotypic variation that could be selected during domestication. Paleologs may be over-represented in our candidate gene lists simply because they contain more genetic diversity than non-paleologs in *B. rapa*. Although many paleopolyploid plant genomes have been analyzed, this difference in genetic diversity has not yet been observed to the best of our knowledge.

The presence of elevated genetic variation in *B. rapa* paleologs suggests that domestication and crop improvement may have proceeded largely from standing variation. Rapid responses to selection, such as the response to selection to domestication and crop improvement, will proceed much faster if standing genetic diversity is high enough to facilitate simultaneous selection at multiple sites (Barrett & Schluter, 2008; Messer & Petrov, 2013; Matuszewski *et al*., 2015). It is an open question how much domestication has proceeded from standing genetic diversity, but a recent analysis suggests that maize could have been domesticated from standing variation in teosinte (Yang *et al*., 2019). There are also examples of candidate genes associated with crops, including in *Brassica oleracea (Purugganan et al., 2000)*, that appear to be selected from standing genetic diversity. Given the elevated diversity of paleologs in *B. rapa*, we may expect domestication in these crops to be dominated by soft sweeps (Messer & Petrov, 2013; Hermisson & Pennings, 2017). A complicating factor in characterizing hard and soft sweeps in *B. rapa* at the moment is the significant difference in genetic variation between paleologs and non-paleologs that may skew such diversity based analyses. More sophisticated approaches that leverage recent advances in deep learning (Schrider & Kern, 2018; Kern & Schrider, 2018; Flagel *et al*., 2019) that are trained to account for differences in variation because of gene origins will likely overcome these issues. Ultimately, this will allow us to characterize the genetics of domestication in the diverse crop varieties of *B. rapa* while providing new insight into how paleopolyploidy may influence the architecture of adaptation in plant genomes.

The ancient hexaploidy in the Brassiceae has been hypothesized to be the source of the outstanding diversity of *Brassica* crop varieties (Lukens *et al*., 2004; Cheng *et al*., 2014, 2016). Our results support this hypothesis by finding that candidate genes for domestication of *B. rapa* crop varieties are enriched in regions of the genome duplicated in the mesohexaploidy. It remains to be seen if paleologs are also significantly enriched in the candidate domestication genes in the crops of other *Brassica* species. Other types of genetic variation, such as structural variants like presence/absence and copy number variants, may also be influenced by post-polyploid genome evolution and play a role in domestication (Golicz *et al*., 2016; Song *et al*., 2020; Gabur *et al*., 2020). Developing pan-genome resources (Bayer *et al*., 2020) for *B. rapa* would allow us to better assess diversity across the crop varieties, and test how different types of genetic diversity have contributed to domestication. *Brassica* are known as the dogs of the plant world for the incredible morphological diversity and number of different domesticated crops compared to other plants (Cheng *et al.,* 2014; Liu *et al*., 2014; An *et al*., 2019). Experimental evolution studies in *B. rapa* have also been able to rapidly select for new pollination syndromes (Gervasi & Schiestl, 2017; Schiestl *et al*., 2018). The higher genetic diversity of paleologs in *Brassica* may play a significant role in this morphological diversity and the rapid responses to selection. However, paleopolyploidy is common among flowering plants with the average species experiencing nearly five rounds of WGD in its ancestry (One Thousand Plant Transcriptomes Initiative, 2019; Li & Barker, 2020). If elevated variation in paleologs is a general phenomenon, then we would expect it to be important in the domestication of many other crops and it may not completely explain the outstanding diversity of *Brassica* crops. Further analyses of paleolog variation and enrichment in other crops are needed to understand the generality of our findings and the implications for the domestication of diverse crops like in *Brassica*. Presently, our results provide a testable hypothesis to explain the observed correlation of polyploidy and crop domestication (Salman-Minkov *et al*., 2016).

More broadly, our results suggest that paleopolyploidy may leave behind a legacy of elevated genetic diversity across the duplicated remnants of diploidized genomes. Although most models and studies of polyploid evolution compare diploids and polyploids (Otto & Whitton, 2000; Ramsey & Schemske, 2002; Otto, 2007; Selmecki *et al*., 2015; Laport & Ng, 2017; Paape *et al*., 2018; Monnahan *et al*., 2019; Han *et al*., 2019; Baniaga *et al*., 2020), our comparison of paleologs and non-paleologs within a diploidized paleopolyploid uncovered evidence for similar dynamics ongoing within plant genomes even millions of years after whole genome duplication. The extensive genome duplication history of plants may result in genomes with different levels of diversity based on the mechanisms of gene origin and retention. Although this remains to be broadly tested, it raises a few testable predictions. If this is a general phenomenon in plants, then we may expect that diploid plants with more paleologs will have higher genetic diversity than those with fewer paleologs. As paleologs are lost over time due to fractionation and gene turnover, we would also expect genetic diversity in diploid plants to be correlated with the time since their most recent paleopolyploid event. The long-term effects of paleopolyploidy on genetic diversity observed here in *B. rapa* may also help explain a broader phenomenon, the lag time of paleopolyploidy and diversification (Schranz *et al*., 2012). Recent research has found that paleopolyploidy in plants is associated with diversification rate increases, but these rate increases often occur many millions of years following WGDs (Landis *et al*., 2018). If relatively high genetic diversity is important for the adaptation and expansion that leads to macroevolutionary signatures of net diversification rate increases, then the observed lag between diversification and polyploidy observed in many studies could be explained by the lasting increase in diversity at paleologs. Although our analyses are centered on how ancient polyploidy contributed to the diversity of *B. rapa*, additional analyses in other plants will provide crucial data to test the new hypotheses described above. Given the distribution of polyploidy throughout the history of flowering plants (One Thousand Plant Transcriptomes Initiative, 2019), our results suggest that the genetic legacy of these WGDs could contribute to the diversity and adaptation of plants millions of years later.

## Acknowledgements

We thank A. Baniaga, G. Finch, Z. Li, and B. Sutherland for feedback and comments on drafts of this manuscript. Hosting infrastructure and services provided by the Biotechnology Computing Facility (BCF) at the University of Arizona. This research was supported by NSF-IOS-1339156 to M.S.B., G.C.C., and J.C.P.

## Author Contributions

H.A. conducted the RNA experiments. X.Q. conducted gene age distribution, candidate gene survey, and all statistics. X.Q. and T.E.H. conducted the differential gene expression analyses. X.Q., C.D., M.T.W.M., and P.D.B. conducted the gene diversity analyses. X.Q. and M.S.B. designed the experiments and wrote the manuscript with input from all co-authors. All authors read and approved the manuscript.

## Data availability

All the data sets generated during the current study are available in the NCBI Sequence Read Archive (SRA) under accession number SRP072186 (http://www.ncbi.nlm.nih.gov/sra/SRP072186). The accession numbers are summarized in Supplementary Table 2.

## Supporting Information

**Fig S1: Manhattan plots of the SweeD analyses.** Each column represents a *B. rapa* genetic group: TO, toria; IS, Indian sarson; PC, pak choi; CC, Chinese cabbage. Each row represents a *B. rapa* chromosome (A01-A10). For each plot, the x-axis denotes the chromosome position (unit: bp), whereas the y-axis denotes the CLR value calculated in SweeD. Each dot denotes the CLR value of a genomic region. Outlier regions were indicated with red dots.

**Fig S2: Circos plot of SweeD (a) and Gene Differential Expression (GDE) gene (b) density across the *B. rapa* genome.** A01-A10 represent the 10 chromosomes of *B. rapa*. The four histogram layers denote the number of identified candidate genes. TO, toria (orange); IS, Indian sarson (light green); CC, Chinese cabbage (olive); PC, pak choi (red). The rainbow ribbons in the center represent the syntenic regions among chromosomes.

**Fig S3: Heatmaps of expression level of genes with significantly different expression in four *B. rapa* crop varieties compared to a control group.** TO = toria, IS = Indian sarson, PC = pak choi, and CC = Chinese cabbage. The European *B. rapa* genetic group was used as control. Each column represents one *B. rapa* accession, whereas each row represents one gene with significantly different expression between the focal group and control group.

**Fig. S4. Comparison of nucleotide diversity (π) across *Brassica rapa* paleologs and non-paleologs from the *B. rapa* only synteny + Ks paleolog inference approach.** The bottom and top of each box represents the first and third quartiles, the band inside each box is the median, and the numbers represent the mean π of each category. Nucleotide diversity is shown for all sites, nonsynonymous sites only, and synonymous sites only for paleologs (purple shades) and non-paleologs (orange shades).

**Fig S5: ANGSD Read mapping.** Nucleotide diversity in paleologs and non-paleologs based on read mapping with STAR and *π* calculated from genotype likelihoods in ANGSD. Box plots on left were filtered for only uniquely mapped reads, whereas the box plots on the right included calculations of *π* based on all read mappings (unfiltered). Paleologs had significantly higher levels of diversity than nonpaleologs for all read mappings (unfiltered t = 48.997, df = 25842, p < 2.2 x 10^-16^ for t-test; W = 238,210,946 and p < 2.2 x 10^-16^ for U-test) and when filtering for only uniquely mapping reads (filtered t = 44.11, df = 26044, and p < 2.2 x 10^-16^ for t-test; W = 232,582,598 and p < 2.2 x 10^-16^ for U-test), demonstrating that our results are unlikely to be affected by read mismapping.

**Table S1: *Brassica rapa* gene IDs for genes identified as paleologs and non-paleologs in our analyses.**

**Table S2: Sample information for the 102 *Brassica rapa* accessions used in this study.**

**Table S3: *Brassica rapa* gene IDs for candidate genes identified by our analyses for each crop variety.** TO = toria, IS = Indian sarson, PC = pak choi, and CC = Chinese cabbage.

**Table S4: Total number of candidate genes identified in our analyses and their distribution on the ten chromosomes of *Brassica rapa*.**

**Table S5: Detailed information of the five *Brassica rapa* domestication candidate genes identified in all three of our genome scan approaches.** Annotation information was obtained from the Brassica Database (brassicadb.org/brad/).

**Table S6: Chi-square test results for POInT paleolog enrichment in candidate gene lists and Mann-Whitney U tests of POInT paleolog nucleotide diversity.**

**Table S7: Summary of the *B. rapa* candidate genes identified from published mapping studies.** Brief description of functions, classical gene names, and *Brassica rapa* gene IDs are given for each candidate gene. Paleolog status is indicated as Y (yes) or N (no). If the candidate gene was also identified in our genome scans, the type of scan is indicated with S (SweeD) or D (Differential gene expression).

