## Supplementary figures and images for "Genes derived from ancient polyploidy have higher genetic diversity and are associated with domestication in *Brassica rapa*"

### Figure S1

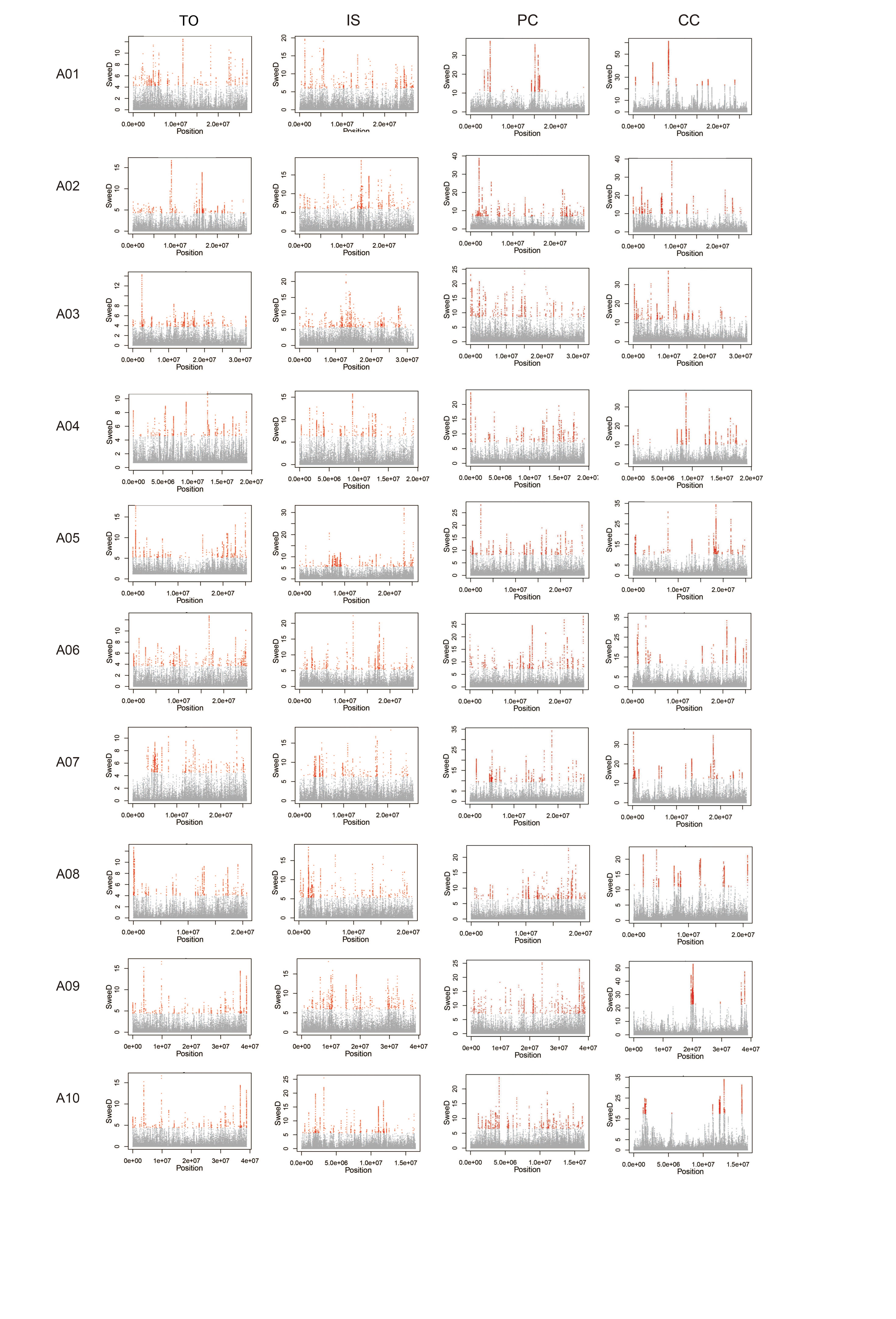

### Figure S2

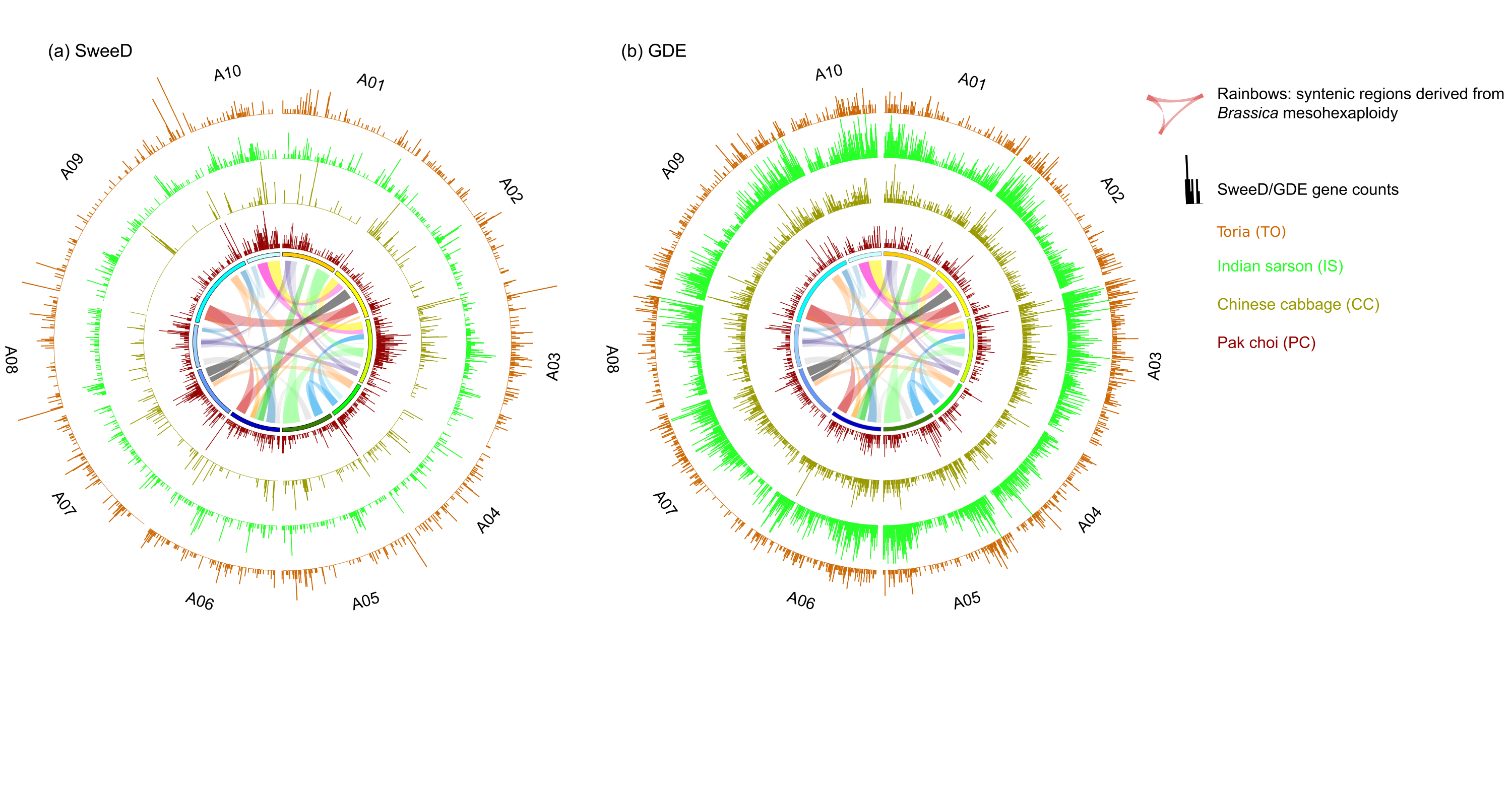

### Figure S3

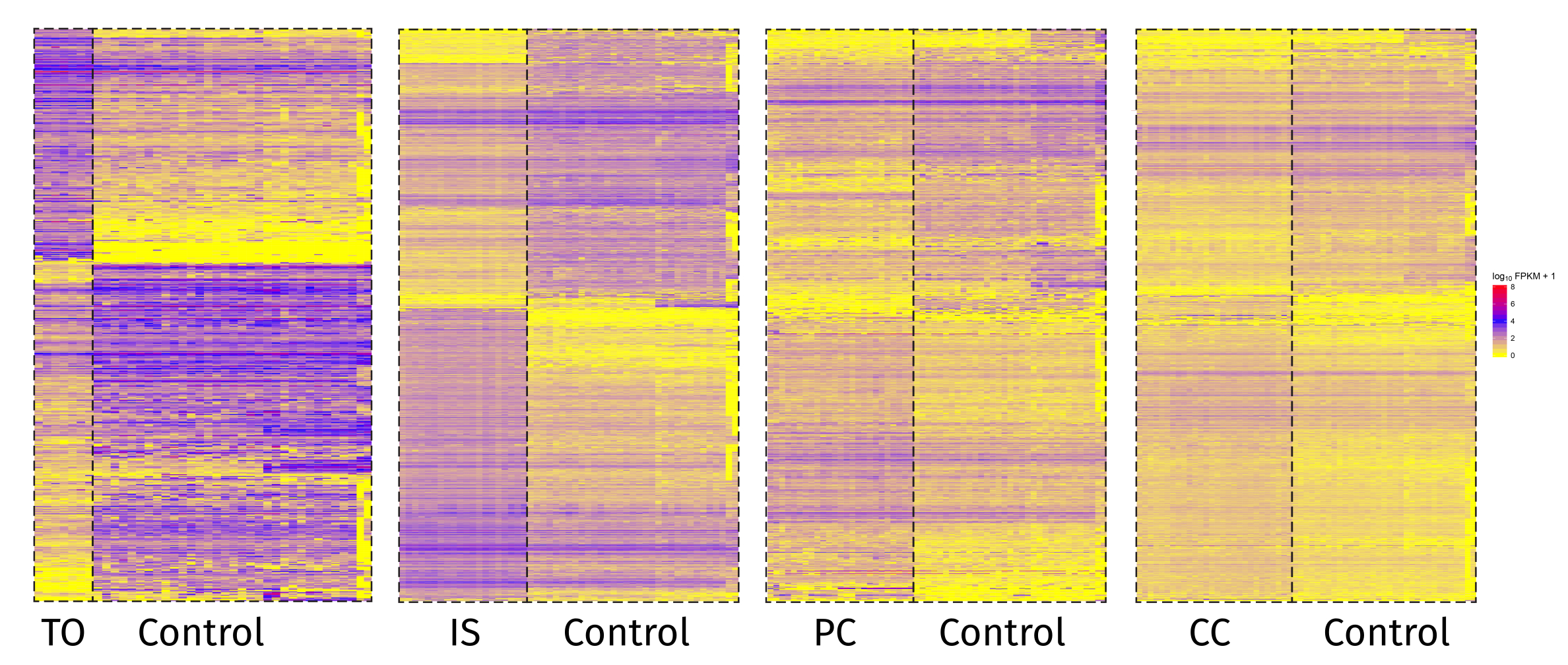

### Figure S4

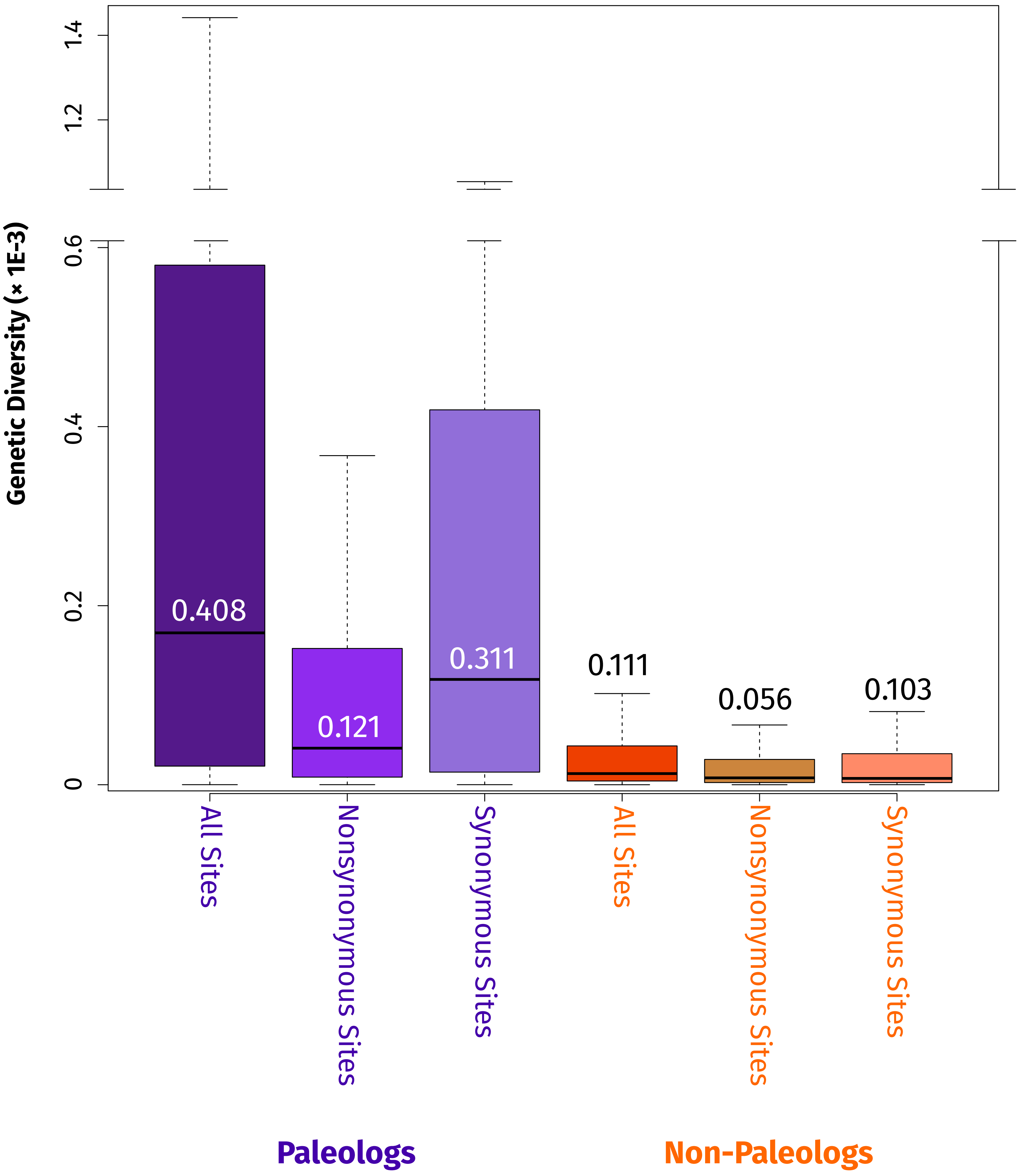

### Figure S5

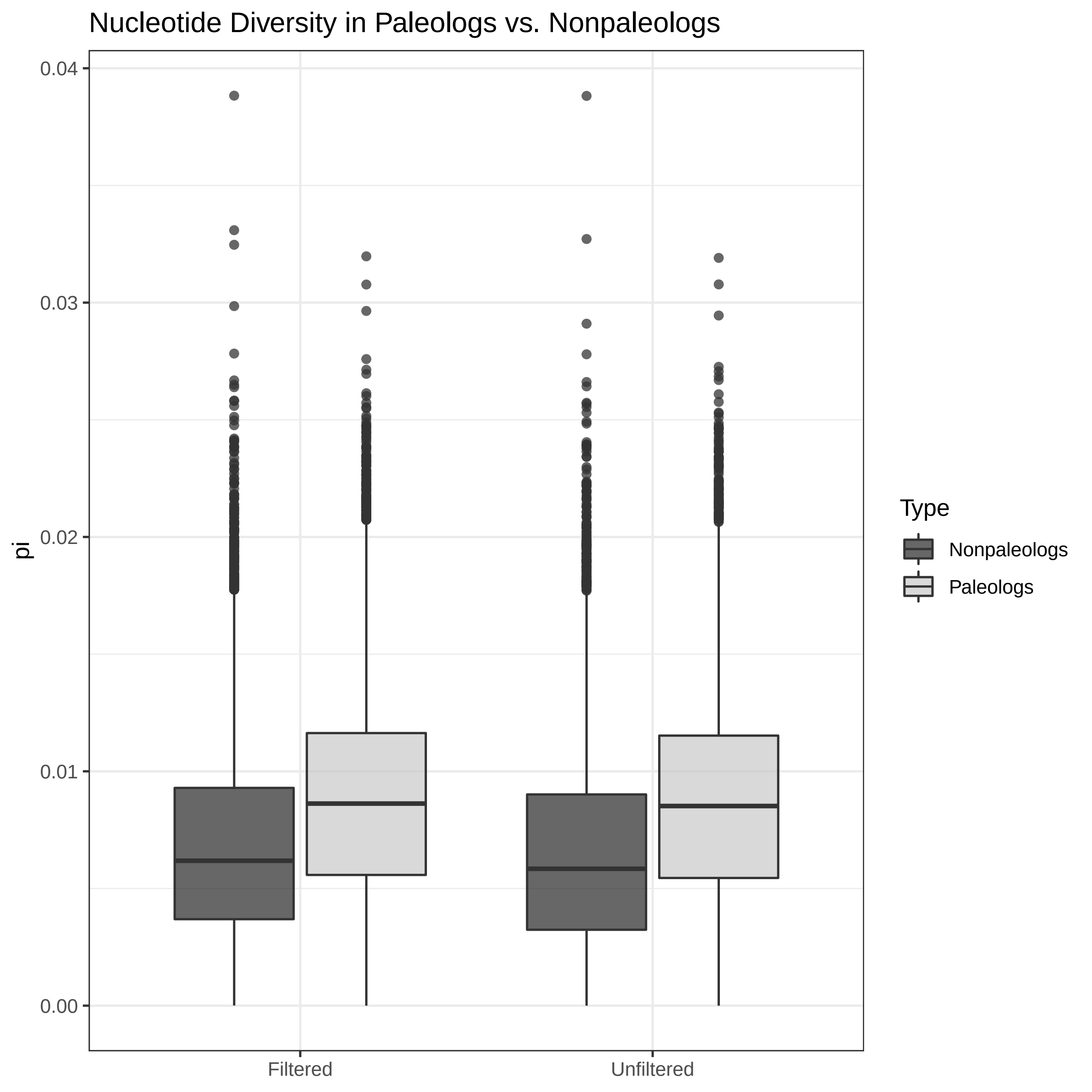
