## Supplementary material for "Genes derived from ancient polyploidy have higher genetic diversity and are associated with domestication in *Brassica rapa*": Table S4

| **Genomic regions under selection** | **SweeD** | **M-K Test** | **DGE** |
| --- | --- | --- | --- |
| **A01** | 300 | 12 | 756 |
| **A02** | 335 | 6 | 774 |
| **A03** | 458 | 8 | 1116 |
| **A04** | 235 | 11 | 488 |
| **A05** | 353 | 9 | 792 |
| **A06** | 338 | 8 | 777 |
| **A07** | 296 | 7 | 710 |
| **A08** | 303 | 6 | 591 |
| **A09** | 447 | 14 | 1064 |
| **A10** | 322 | 9 | 555 |
| **Scaffolds** |  | 2 | 190 |
| **Total** | **3387** | **92** | **7813** |
