## Supplementary material for "Genes derived from ancient polyploidy have higher genetic diversity and are associated with domestication in *Brassica rapa*": Table S5

| **Gene accession** | **Chromosome** | **Position** | **Length** | **CDS number** | **Related *Arabidopsis* gene** | **GO annotation** | **KEGG annotation** | **Swissport annotation** |
| --- | --- | --- | --- | --- | --- | --- | --- | --- |
| ***Bra010933*** | A08 | 16417093  -16420284 | 3192 | 11 | AT1G27510 | N/A | hypothetical protein ; F-box protein 31 | EXEC1_ARATH Protein EXECUTER 1 |
| ***Bra023190*** | A09 | 20936190-20941067 | 4878 | 21 | AT1G31480 | metal ion binding; | DDHD1; acetyl esterase | DDHD1_HUMAN Phospholipase DDHD1 |
| ***Bra031452*** | A01 | 17166656-17171527 | 4872 | 11 | AT1G61140 | zinc ion binding; ATP binding;  protein binding; helicase activity;  DNA binding;  nucleic acid binding; | RAD16; nucleotide excision repair protein; adenosinetriphosphatase | Putative SWI/SNF-related matrix-associated actin-dependent regulator of chromatin subfamily A member 3-like 2 |
| ***Bra003055*** | A10 | 5699614-5702647 | 3034 | 13 |  | protein amino acid phosphorylation；ATP binding; protein serine/threonine kinase activity; protein kinase activity; | - | - |
| ***Bra035693*** | A04 | 12878256-12883485 | 5230 | 18 | AT2G28520 | proton-transporting two-sector ATPase complex, proton-transporting domain; ATP synthesis coupled proton transport；hydrogen ion transmembrane transporter activity； | K02154; VHA-A1; VHA-A1；ATPase; V-type H+-transporting ATPase subunit | VPP1_XENLA V-type proton ATPase |
