## Supplementary material for "Genes derived from ancient polyploidy have higher genetic diversity and are associated with domestication in *Brassica rapa*": Table S7

**Table S6**

| **Trait** | **Classical Gene / QTL Name** | ***B. rapa* Gene ID** | **Paleolog by *B. rapa* only Method** | **Paleolog by POInT Multi-Genome Method** | **Candidate Gene Scan** | **Reference** |
| --- | --- | --- | --- | --- | --- | --- |
| Anthocyanin pigmentation | *bHLH49* | Bra004348 | Y | N |  | ^1^ |
| Seed coat color | *BrTT1 (Brsc1)* | Bra028067 | Y | N |  | ^2^ |
| Hairiness and seed coat color | *BrTTG1** | Bra009770  Bra029411 | Y  N | Y  Y |  | ^3^ |
| Orange head of Chinese cabbage | *BrOr, BrCRTISO* | Bra031539 | Y | Y | D | ^4–7^ |
| Orange flower of Chinese cabbage | *BrOF* | Bra037124 | Y | N |  | ^8^ |
|  |  | Bra037125 | Y | Y |  |  |
| Clubroot resistance | *Rcr1* | Bra019409 | N | N | D | ^9^ |
|  |  | Bra019410 | Y | N | SD |  |
| Clubroot resistance | *Rcr5* | Bra012595 | Y | N |  | ^10^ |
|  |  | Bra012591 | N | Y |  |  |
|  |  | Bra012588 | Y | Y |  |  |
|  |  | Bra012586 | Y | N | D |  |
|  |  | Bra012581 | Y | Y | D |  |
|  |  | Bra012580 | Y | Y | D |  |
|  |  | Bra012578 | Y | Y |  |  |
| Clubroot resistance | *Rcr6* | Bra010663 | Y | N |  | ^11^ |
| Clubroot resistance | *Rcr9* | Bra020936 | N | N |  | ^12^ |
|  |  | Bra020861 | Y | N |  |  |
| Circadian period | *PERA9a/GI* | Bra024536 | Y | N | D | ^13^ |
| Cuticular wax biosynthesis | *BrWax1* | Bra013809 | Y | N |  | ^14^ |
| Cuticular wax biosynthesis | *BrCER1* | Bra032670 | N | Y | SD | ^15^ |
| Cuticular wax biosynthesis | *BrCER4* | Bra011470 | Y | N | SD | ^16^ |
| Flower organ development | *pdm* | Bra040093 | Y | Y | D | ^17^ |
| Leaf color variation | *lcm1* | Bra006208 | Y | Y | SD | ^18^ |
| Leaf color variation | *pem* | Bra024218 | Y | Y | D | ^19^ |
| Leaf/lobe and hairiness | *BrGA20OX3* | Bra010064  Bra009285  Bra028706  Bra005927 | N  Y  Y  Y | N  N  N  N | D | ^20^ |
| Leaf/heading | *BrARF3.1*  *BrARF4.1*  *BrKAN2.1*  *BrKAN2.3*  *BrBRX.1*  *BrBRX.2* | Bra005465  Bra002479  Bra023254  Bra033844  Bra023219  Bra035521 | Y  Y  Y  Y  Y  Y | Y  Y  N  N  N  N | D  S | ^21^ |
| Stay-green leaves | *Brnye1* | Bra019346 | Y | N |  | ^22^ |
| Tuberous/sugar transporter 1 | *BrSTP1.1*  *BrSTP1.3* | Bra019870  Bra031762 | Y  Y | Y  Y |  | ^21^ |
| Trichomes | *Brtri1** | Bra025311 | Y | Y |  | ^23^ |
